# Structural basis of a phosphotransferase system xylose transporter

**DOI:** 10.1101/2025.04.02.646771

**Authors:** Yutaro S. Takahashi, Hidetaka Kohga, Min Fey Chek, Kotomi Yamamoto, Jun F. Takahashi, Hideki Shigematsu, Yoshiki Tanaka, Muneyoshi Ichikawa, Ryoji Miyazaki, Toshio Hakoshima, Tomoya Tsukazaki

## Abstract

The bacterial phosphotransferase system (PTS) is essential for carbohydrate uptake, and IIC transporters in the system facilitate intracellular sugar transport. Here, we report the crystal and cryo-EM structures of *Leminorella grimontii* GatC (LgGatC), a putative xylose transporter in the IIC family, in its outward-facing state. Structural analysis indicates that the outward-facing cavity recognizes the linear form of D-xylose. A structure-based homology model of an inward-facing state supports an elevator-like sugar transport mechanism. These findings provide insights into PTS-mediated xylose transport and its potential applications in microbial engineering.

## Introduction

Xylose, a major component of hemicellulose, constitutes 18–30% of sugars released when lignocellulose is hydrolyzed^1^. Although xylose is considered a promising resource for the microbial production of biobased compounds such as lactate^2–5^, its utilization as a carbon source faces significant challenges. These challenges occur because only a limited number of microorganisms possess xylose metabolic pathways, and even those pathways capable of its metabolism are often restricted by glucose-induced carbon catabolite repression (CCR)^6^; this further complicates its efficient use. The uptake of xylose across the biological membrane is the first step in its cellular utilization and significantly influences both the conversion efficiency and the cell growth rate^7,8^. Therefore, a detailed understanding of the xylose uptake mechanism can likely facilitate the efficient production of versatile compounds using xylose as a raw material. Studies on *Escherichia coli* strains that ferment xylose have indicated that the phosphoenolpyruvate-dependent phosphotransferase system (PTS) enzyme IIC component (IIC) GatC facilitates intracellular xylose uptake^9^. These results are further supported by adaptive laboratory evolution experiments in *E. coli*, which were designed to increase xylose fermentation efficiency^3^.

The PTS consists of cytoplasmic proteins, including histidine phosphocarrier protein (HPr) and enzyme I (EI), as well as substrate-specific II factors (IIA, IIB, and IIC) (Fig. S1a). Intracellular phosphoenolpyruvate (PEP) serves as a phosphate donor and initiates a stepwise phosphorylation process. Ultimately, through the coordinated action of IIB and IIC, sugars are transported into the cell in a phosphorylated form. PTS systems are classified into four evolutionarily distinct superfamilies: (i) glucose-fructose-lactose (GFL), (ii) ascorbate-galactitol (AG), (iii) mannose-PTS, and (iv) dihydroxyacetone (DHA)^10–12^. The putative xylose transporter GatC belongs to the AG superfamily.

Crystal and cryo-EM structures of the IIC proteins belonging to the GFL, AG, and mannose-PTS superfamilies have been reported^13–19^. IIC proteins in the GFL and AG superfamilies typically form homodimers (Fig. S1b, c, d), whereas the mannose-PTS IIC uniquely associates with IID proteins, forming IIC-IID heterodimers that further assemble into functional trimers^18^ Despite the structural differences in folding between the GFL and AG superfamilies, their IIC proteins share a core structure consisting of an interaction domain (scaffold or V motif domain) and a substrate-binding domain (transport or core domain) (Fig. S1b, c, d). IIC proteins facilitate substrate transport through structural transitions from an outward-facing to an inward-facing state, accompanied by rigid rotation of the substrate-binding domain. The L-ascorbic acid-specific IIC protein UlaA, which is a member of the AG superfamily, including GatC, has been structurally characterized using *E. coli* and *Pasteurella multocida* UlaA^16,17^ and shares approximately 12% sequence identity with *Leminorella grimontii* GatC (LgGatC), which is the structural target in this study. The detailed structure of GatC will provide critical insights for the rational design of the engineered variants with improved transport functionality, advancing the development of a xylose-based bioproduction platform. This study presents the crystal and cryo-EM structures of LgGatC with/without xylose in an outward-facing form. These structures and the proposed homology model of LgGatC in an inward-facing state provide crucial insights into the mechanism of GatC-mediated xylose transport.

## Results and Discussion

### Functional characterization and crystal structure of LgGatC

To identify a suitable target for structural analysis of GatC, we first assessed its stability and accumulation levels using fluorescence-detection size exclusion chromatography (FSEC) screening^20^ of GatC from various species, including *E. coli* (Ec) and *L. grimontii* (Lg). Each full-length GatC tagged with a C-terminal GFP-His_8_ was expressed in *E. coli* cells, solubilized from the membrane fraction using n-dodecyl-β-D-maltoside (DDM), and analyzed via the FSEC system^20^. Since LgGatC presented the highest monodispersity and accumulation levels, it was selected for further examination. LgGatC shares 81% sequence identity with EcGatC (Fig. S2), indicating functional similarity; however, its substrate remains unknown. To investigate whether LgGatC binds D-xylose, we performed isothermal titration calorimetry (ITC) experiments, considering the proposed D-xylose transport activity of EcGatC^3,9^. The dissociation constant (K_d_) between LgGatC and D-xylose was approximately 48.8 µM; these results indicate that LgGatC, similar to EcGatC, functions as a D-xylose transporter (Fig. 1a).

**Figure 1.**
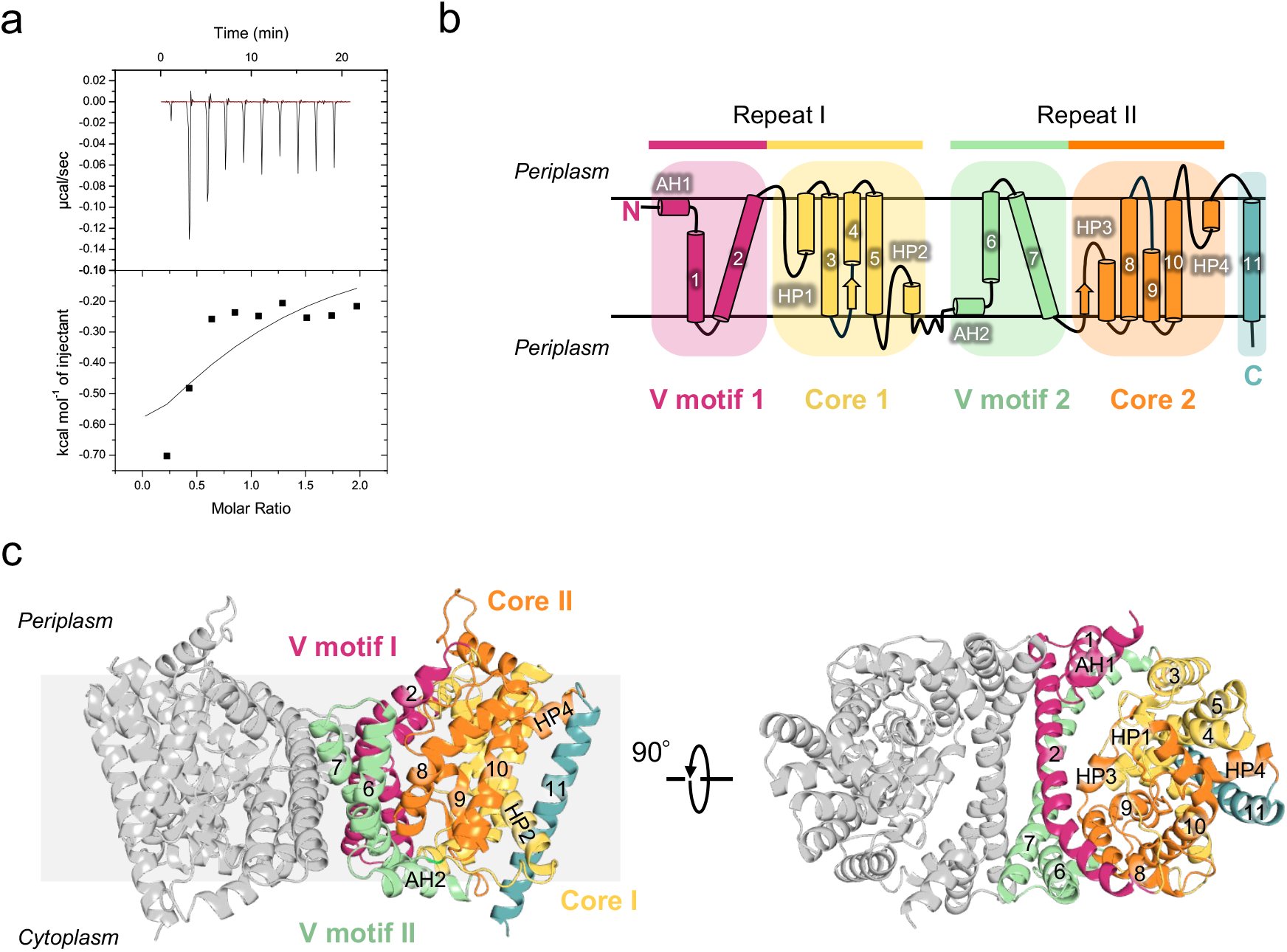
Functional and structural characterization of *Leminorella grimontii* GatC. (a) ITC data of LgGatC binding to D-xylose. The top panel shows the raw heat changes. The bottom panel presents the binding isotherm. Curve fitting of the isotherms reveals that the Kd, ΔH and ΔS values are 48.8 μM, -1141± 513.7 cal/mol, and 15.8 cal/mol/deg, respectively. (b) Topology and domains of the LgGatC protomer. The LgGatC protomer consists of V motif 1 and Core 1 in the N-terminal region, followed by a similar repeated structure of V motif 2 and Core 2. At the C-terminal end, a non-repeated TM11 is present. AH: amphipathic α-helices. HP: α-helical hairpin-like structure. (c) Crystal structure of dimeric LgGatC in the outward-facing form. One protomer is shown in gray, while the other protomer is colored as shown in (b).

LgGatC was crystallized using the lipidic cubic phase (LCP) method in the presence and absence of xylose (10 mM and 0 mM, respectively). The crystal structures of the LgGatC homodimer under both conditions (excluding each C-terminal region from residue 449) were determined by X-ray crystallography at resolutions of 3.30 Å and 3.74 Å, respectively (Fig. 1c; Table S1). A sequence comparison between LgGatC and *Pasteurella multocida* (Pm) UlaA, an AG superfamily IIC protein, suggested that the topologies are similar and that the C-terminus of LgGatC is oriented toward the cytoplasm (Fig. S3). The structures of LgGatC in the presence and absence of xylose are highly similar, and the superposition of the overall dimer structures yields a root mean square deviation (RMSD) of 0.168 Å for all Cα atoms (Fig. S4). Each LgGatC protomer consists of 11 transmembrane helices (TMs), four reentrant α-helical hairpin-like structures (HP1– 4), and two horizontal amphipathic α-helices (AH1 and AH2) (Figs. 1b, c, S2). Except for TM11, the protomer contains an internal inverted structural repeat, with Repeat I (N-terminal) symmetrically inverted relative to Repeat II (C-terminal) in the membrane (Fig. 1b). Each initial region in Repeats I and II consists of AH1–TM1–TM2 and AH2–TM6– TM7, respectively, and forms V-shaped motifs, which are referred to as V motif 1 and V motif 2. The TM11, the final transmembrane helix, is positioned at the C-terminal end. The LgGatC structures exhibit an outward-facing state, as the core domain contains a cavity accessible from the periplasmic side.

### Substrate recognition mode and Cryo-EM analysis of LgGatC

Residues W83 and S87 on HP1; N125, W127, and H128 on TM4; G182, T183, and S184 on HP2; D296 and P297 on HP3; F340 on TM9; and D399 on HP4 form the negatively-charged cavity in the core domain (Fig. S5). The X-ray crystallographic electron density map shows smaller blobs than the xylose molecules (Fig. S5). Although the blobs observed in the presence of xylose were larger than those in its absence, we were not able to determine whether these blobs originated from xylose. To obtain further structural insights, LgGatC-reconstituted nanodiscs^21^ with *E. coli* lipids were analyzed using cryo-EM. The global resolution of the 3D maps generated from single-particle analysis was 2.70 Å for the xylose-bound state and 3.27 Å for the apo state (Figs. 2a, b, S6, 7). The two cryo-EM structures of LgGatC correspond to the same outward-facing state as the crystal structure with the negatively charged cavity (Figs. 2a–d). A superposition comparison of the crystal and cryo-EM structures, which was based on Cα atoms, revealed an RMSD of 0.663 Å between the apo-state and D-xylose-bound cryo-EM structures and 0.889 Å between the crystal and cryo-EM structures in the presence of D-xylose (Fig. S4). These low RMSD values indicated that all the structures determined in this study adopted essentially the same overall conformation; thus, no significant conformational changes occurred regardless of the presence or absence of D-xylose (Fig. S4). Nonprotein Coulomb potential map within the cavity of the cryo-EM structures in the presence of xylose shows a linear shape, whereas a spherical shape was observed in its absence (Fig. 2 e, f). We assigned the linear-shaped density observed in the xylose-bound map to D-xylose (Fig. 2e). In contrast, non-proteinaceous densities identified in the crystal structures and the cryo-EM structure under apo conditions likely originated from water and/or glycerol molecules used during LgGatC purification (Figs. 2f, S5).

**Figure 2.**
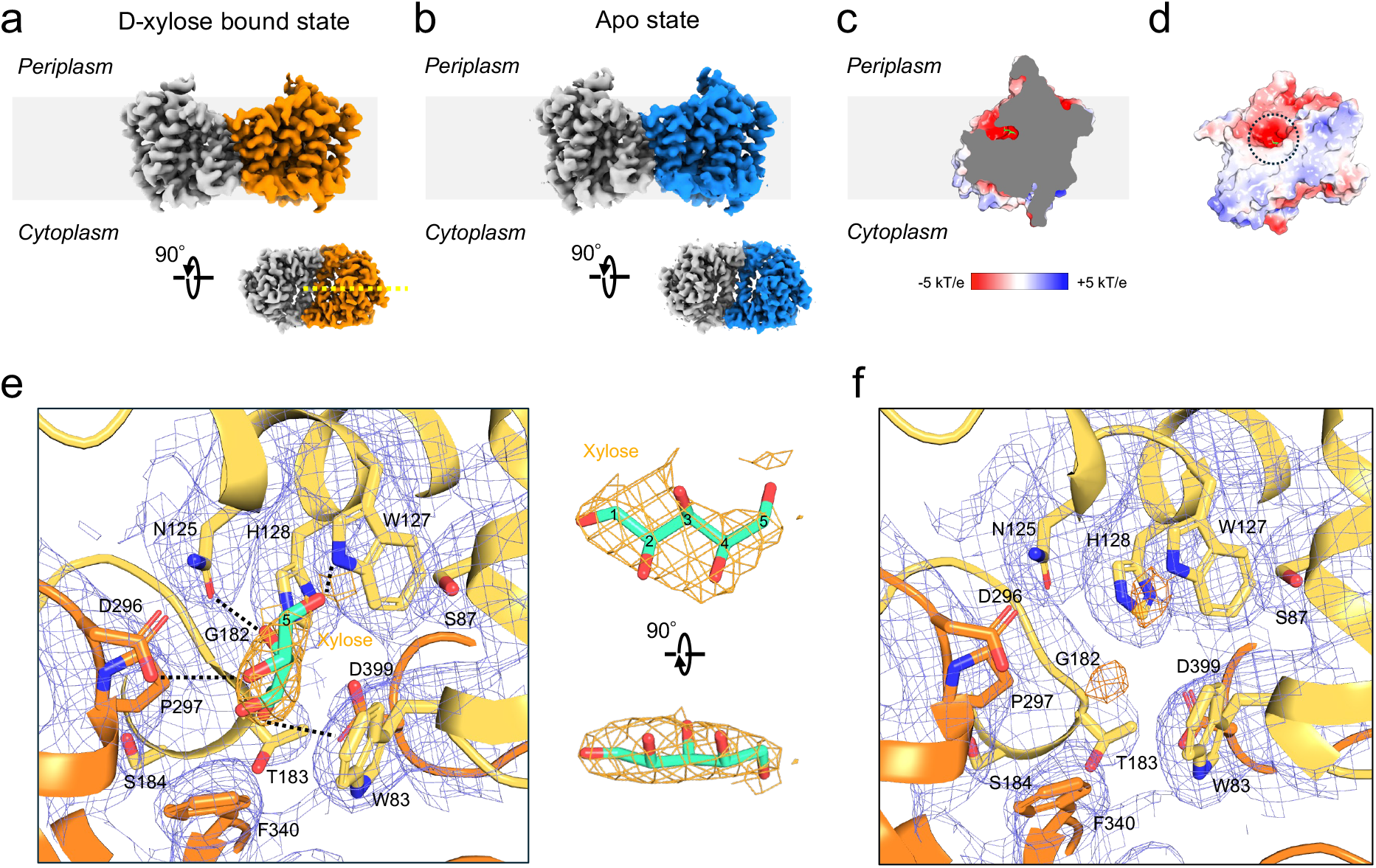
Cryo-EM structures of outward-facing LgGatC. (a) Cryo-EM map of LgGatC in the D-xylose-bound state, contoured at a threshold level of 0.04. One protomer is shown in gray, while the other is colored in orange. (b) Cryo-EM map of LgGatC in the apo state, contoured at a threshold level of 0.07. One protomer is shown in gray, while the other is colored in blue. The nanodisc Coulomb potential map is omitted for clarity. (c) Negatively charged cavity of LgGatC. Cross-sectional view along the dashed line in (a), showing the electrostatic surface potential ranging from -5 kT/e (red) to +5 kT/e (blue). The D-xylose molecule is shown as sticks. (d) The electrostatic surface of GatC protomer viewed from the dimer interface. (e), (f) Cryo-EM maps in the presence (e, contoured at 5.0σ) and absence (f, contoured at 8.0σ) of D-xylose, shown with the structural cartoon models around the cavity. The side chains of the residues forming the cavity surface are displayed as sticks. The maps are displayed, with the entire map shown in blue and the nonprotein Coulomb potential highlighted in orange. The cryo-EM map corresponding to xylose was shown (e, right). Dashed lines represent potential hydrogen bond interactions. The carbon numbers are shown on the sugar model.

Glycerol functions as a sugar mimetic in the active sites of carbohydrate-processing enzymes^22,23^. D-xylose, a pentose sugar, exists in equilibrium with its pyranose, furanose, and linear forms in solution. Although the physiological substrate of GatC remains unclear, we hypothesize that GatC recognizes the open-chain (liner) form of D-xylose. This hypothesis is supported by reports that GatC transports linearly structured sugar alcohols, such as galactitol, sorbitol (glucitol), and xylose^3,24,25^. Since IIC transporters are proposed to release substrates into the cytoplasm via phosphate transfer in coordination with IIB, the C-5 carbon of D-xylose, which is the site of phosphate modification, was positioned near the cavity entrance in our model (Fig. 2e). Polar residues N125, W127, H128, D296, and D399 in the cavity may form hydrogen bonds with the -OH groups of D-xylose and potentially contribute to the specificity and stabilization. Additionally, the hydrophobic residues W83 and F340 may interact with the carbon chain of D-xylose, further contributing to binding stabilization through hydrophobic interactions (Fig. 2e, f). These residues are highly conserved across GatC orthologs, highlighting their importance in xylose transport (Fig. S2).

### Homology model construction and working model

The crystal structures of EcUlaA and PmUlaA reveal that they exist in outward-facing, occluded, and inward-facing states^16,17^; this leads to the prediction that UlaA transports the substrates into the cell through an elevator-type conformational change involving rigid body movement of the core domain^16,17^ The LgGatC structures in this study were compared with those of EcUlaA and PmUlaA and revealed that GatC lacks an AH region in UlaAs connecting TM6 and TM7 and shares only approximately 12% sequence identity with UlaA (Fig. S3). However, since their overall secondary structural features are identical except for the AH, GatC is likely to employ the same molecular transport mechanism as UlaA. TM2 and TM7 of the V-motif subdomain in the UlaA structure form a curved structure in which glycine residues G58 and G286 serve as pivot points that facilitate the rigid body movement of the core domain during substrate transport^26^. Similar to UlaA, LgGatC has highly conserved glycine residues G49 and G270 on TM2 and TM7 (Fig. S2, S3), which potentially contribute to the mobility of the V-motif domain. To gain further insight into the substrate transport mechanism of LgGatC, we generated an inward-facing homology model on a SWISS-MODEL^27^ server (Fig. 3a) by uploading the structure-based amino acid alignment between GatC and UlaA (Fig. S3) using the inward-facing structure of PmUlaA (PDB: 5ZOV) as a template. In the outward-facing crystal structure of LgGatC, residues W83, N125, H128, G182, T183, D296, F340, and D399 contribute to cavity formation (Fig. 2d), whereas in the inward-facing homology model, these residues are repositioned near the cavity region rather than within the cavity itself. Additionally, based on the predictions from the positioning of proteins in membranes (PPM) server^28^, the core domain could undergo a rigid body rotation, whereas the transmembrane region of the V-motif domain remains unchanged between the outward-facing structure and the inward-facing homology model (Fig. 3b). This structural comparison suggests that the GatC core domain undergoes a rigid conformational shift across the plasma membrane during the transition from the outward-facing to the inward-facing state, resembling the proposed structural mechanism of UlaA. In this study, the structural characterization of LgGatC as a PTS galactitol family protein IIC was presented and supported a model in which a transport mechanism similar to the UlaA elevator model drives substrate movement in GatC (Fig. 3c). In this model, GatC in the outward-facing form selectively binds D-xylose on the periplasmic side. Upon substrate binding, the core domain undergoes a rigid-body rotational conformational transition to the inward-facing form, transporting D-xylose across the membrane. Finally, GatC releases the substrate into the cytosol.

**Figure 3.**
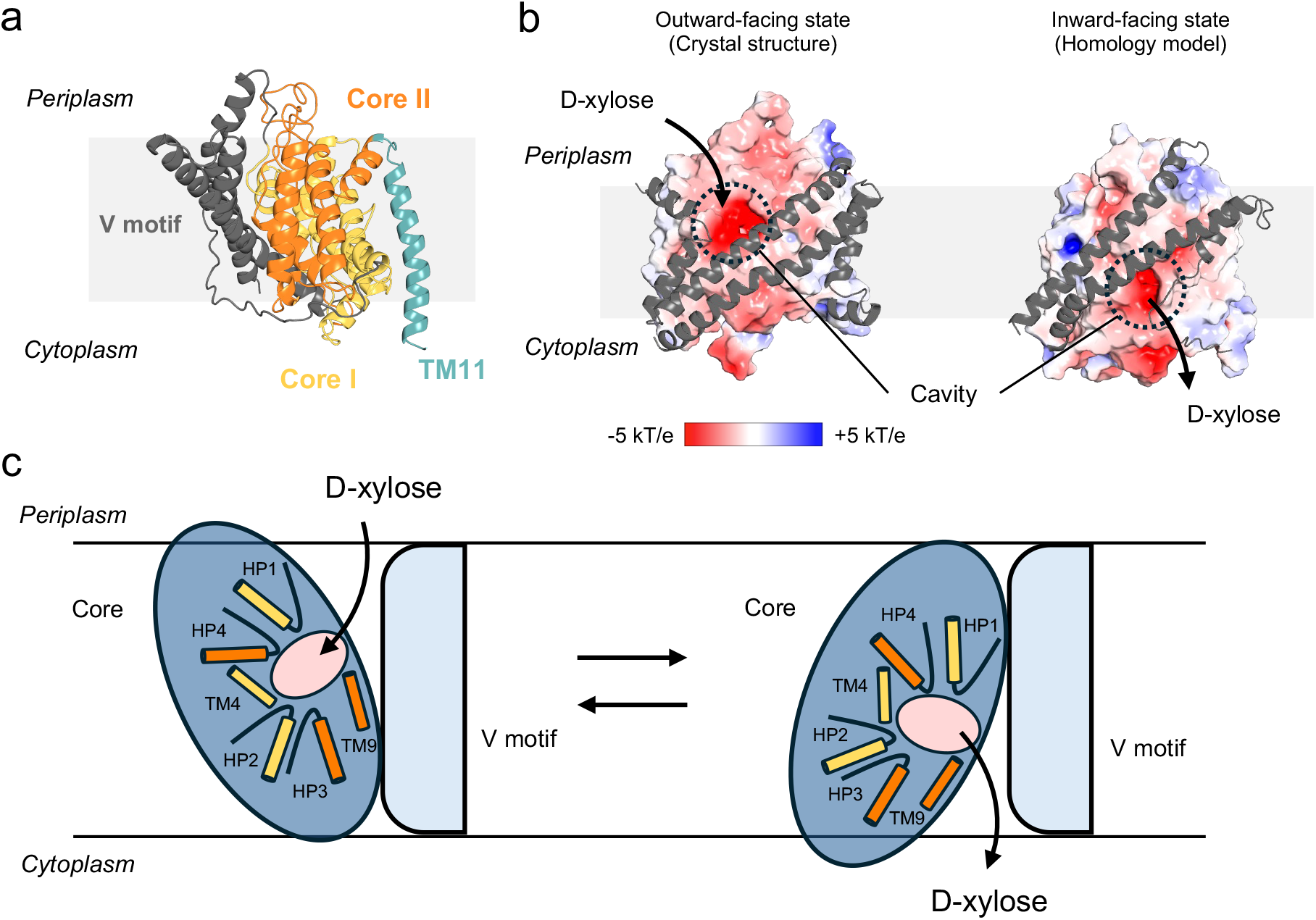
Working model of alternating outward- and inward-facing conformations of LgGatC in D-xylose transport. (a) Homology model of inward-facing LgGatC prepared using the amino acid alignment based on the structural superimposition of LgGatC and EcUlaA (PDB ID: 4RP9) (Fig. S3). The V-motif domain is shown in gray, and the remaining regions are colored as in Fig. 1. (b). The spatial position of the homology model inside the lipid bilayer was predicted by the PPM server. (b) Comparison of the core domain between the outward-facing crystal structure and the inward-facing homology model. The V-motif domains are shown in gray ribbons, whereas the remaining regions are displayed by electrostatic potential surfaces ranging from -5 kT/e (red) to +5 kT/e (blue). (c) D-xylose transport working model of GatC. First, D-xylose is recognized by the outward-facing state. Then, a conformational change to the inward state, with the core domain acting as a rigid body, releases the substrate into the cytoplasm.

## Acknowledgments

We thank Kayo Abe for secretarial assistance, Kunihito Yoshikaie for technical support, and the scientists of SPring-8 Structural Biology Beamlines for helping with data collection. This work was supported by JSPS/MEXT KAKENHI (Grant No. JP23K14146 to H.K., Grant Nos. JP21K15028 to M.F.C.; Grant Nos. JP22K15061, JP22H05567 to R.M.; and Grant Nos. JP22H02567, JP22H02586, JP21H05155, JP21H05153, JP21KK0125 to T.T.); PRESTO (JPMJPR20E1 to M.I.); Natural Science Foundation of Shanghai (24ZR1403800 to M.I.); private research foundations (Yanmar Environmental Sustainability Support Association to Y.S.T; The Chemo-Sero-Therapeutic Research Institute to T.T.; and Takeda Science Foundation to T.H. and T.T.); and JST SPRING (Grant No. JPMJSP2140 to Y.S.T and J.F.T). The synchrotron radiation and cryo-EM experiments were performed at SPring-8 with the approval of the Japan Synchrotron Radiation Research Institute (JASRI) (Proposal Nos. 2021A2745, 2022A2738, 2023A2727, 2024A2742, 2024A2759). This research was partially supported by Platform Project for Supporting Drug Discovery and Life Science Research (Basis for Supporting Innovative Drug Discovery and Life Science Research (BINDS)) from AMED under Grant Number JP24ama121001.

## Author contributions

T.H. and T.T. conceptualized the study. Y.S.T. and K.Y. purified the samples. Y.S.T., K.H., K.Y., J.F.T., H.S., Y.T., M.I., R.M., and T.T. determined the structures. M.F.C. and T.H. performed ITC experiments. Y.S.T. and T.T. wrote the manuscript. T.T. supervised the study.

## Data availability

The atomic coordinates and structural factors for the crystal structures have been deposited in the Protein Data Bank (PDB) under accession codes 9U8E and 9U8H. The cryo-EM maps and the corresponding atomic models have been deposited in the EM Data Bank (EMDB) under accession codes EMD-63950 and EMD-63951 and in the PDB under accession codes 9U82 and 9U84, respectively.

## Materials and methods

### Purification

The *E. coli* strain C41 (DE3) (Lucigen) containing a modified pCGFP-BC^20^ vector that encodes *Lg*GatC (NCBI Reference Sequence: WP_261832779.1)-LESSGENLYFQFTSSV-GFP_2-238_-His_8_ or -LESSGENLYFQFTSSV-His_8_ was cultured in LB broth Lennox (Nacalai) supplemented with ampicillin (50 µg/mL) at 37°C. When the OD_600nm_ reached 0.6, protein expression was induced with 1 mM isopropyl *β*-D-thiogalactopyranoside (IPTG) at 20°C for 18 h. Harvested cells were lysed at 100 MPa in 300 mM NaCl, 50 mM Tris-HCl (pH 8.0), and 1 mM phenylmethylsulfonyl fluoride (PMSF) using a microfluidizer M-110EH (Microfluidics). The lysate was centrifuged at 8,000 rpm for 30 min (Himac R13A rotor), and the supernatant was ultracentrifuged at 40,000 rpm for 60 min (Beckman 45Ti rotor). The precipitated membrane fraction was resuspended in Buffer A [300 mM NaCl, 50 mM HEPES-NaOH (pH 7.0), 10% glycerol] containing 2% (w/v) DDM (Glycon) and 5 mM β-mercaptoethanol (β-ME). The solution was rotated at 4°C for 60 min. After ultracentrifugation (40,000 rpm for 30 min, Beckman 70Ti rotor), the supernatant was mixed with TALON resin (Clontech) preequilibrated with Buffer A containing 0.1% DDM and 20 mM imidazole-HCl (pH 7.0) for 1 h. The resin was washed with Buffer A containing 0.1% DDM and 20 mM imidazole-HCl (pH 7.0), and then the protein was eluted with Buffer A containing 0.1% DDM and 200 mM imidazole-HCl (pH 7.0). Fractions containing the target protein were pooled. The GFP-His tag or His tag was cleaved by TEV (S219V) protease^29^ cleavage at 4°C for 16 h. The released tag and TEV protease were removed using Ni-NTA agarose resin (QIAGEN) preequilibrated with Buffer B (150 mM NaCl, 50 mM HEPES-NaOH (pH 7.0), 0.05% DDM), and the flow-through fraction was concentrated with an Amicon Ultra 50K NMWL (Merck Millipore). The concentrated sample was further purified using size exclusion chromatography on a Superdex 200 Increase 10/300 (GE Healthcare) column equilibrated with Buffer B. The fraction containing *Lg*GatC was concentrated to 20 mg/mL using an Amicon Ultra 50K NMWL for crystallization and stored at -80°C until use.

### Isothermal titration calorimetry

Purified LgGatC protein (∼20 mg/mL) was dialyzed overnight at 4°C against ITC buffer (150 mM NaCl, 50 mM HEPES-NaOH pH 7.0, 0.05% DDM). The dialyzed GatC sample was collected and centrifuged at 13,000 rpm for 10 min (TOMY AR015-24 rotor). The GatC protein concentration was adjusted to 50 μM using the ITC buffer (dialysate). D-xylose was dissolved in the ITC buffer to prepare a 500 μM solution. The ITC experiment was performed at 20°C using a MicroCal iTC200 (GE Healthcare). To account for the heat of dilution, a 500 μM D-xylose solution was titrated against the ITC buffer. The data were analyzed using Origin 7 software (OriginLab).

### Crystallization

The purified *Lg*GatC protein solution was mixed with monoolein (NU-CHEK-PREP) at a 2:3 protein-to-lipid ratio (v/v) for LCP reconstitution using the twin-syringe method^30^. Each 30 nL aliquot of the mixture was dispensed onto Medical Research Council under-oil crystallization plates (Hampton Research) and overlaid with 3 µL of a reservoir solution, which consisted of 100 mM Na-acetate, 100 mM Tris-HCl (pH 8.0), 40–45% polyethylene glycol (PEG), and 0 or 10 mM D-xylose, using the gryphon protein crystallization system (Art Robbins Instruments). The crystals of LgGatC were grown at 20°C over 4-7 days. The crystals were directly harvested from the drops, flash-cooled in liquid nitrogen, and stored in liquid nitrogen until X-ray diffraction analysis.

### Crystal structure determination of *Lg*GatC

X-ray diffraction experiments were conducted at beamline BL32XU of SPring-8, and the datasets were obtained through the automated ZOO^31^ data collection system. The complete datasets were generated by merging multiple small-wedge datasets collected from hundreds of microcrystals, each approximately 20 μm in size. Data processing was performed using kernel application for the multi-crystal data optimization (KAMO)^32^ on XDS^33^. The initial phase of the LgGatC dataset was determined by molecular replacement using PHASER^34^ with an AlphaFold (LocalColabFold ver. 1.3.0)^35,36^ prediction model as a template. The LgGatC structure in the presence of D-xylose was manually fitted to the density using COOT^37^. Iterative refinement and model modification were performed using PHENIX^38^ and COOT^37^ until R_work_/R_free_ converged to 0.2223/0.2786 at a resolution of 3.3 Å. The structure of LgGatC in the absence of D-xylose was similarly determined by molecular replacement and refined using the same strategy, resulting in final R_work_/R_free_ values of 0.2536/0.2855 at 3.74 Å resolution. The structural figures were generated using PyMOL (https://pymol.org/2/).

### Reconstitution of LgGatC into lipid nanodiscs

Assembly of GatC-containing nanodiscs: *E. coli* total lipid extract (Avanti) was dissolved in 20 mM HEPES-NaOH (pH 7.0), 150 mM NaCl, 5% glycerol, and 0.1% DDM. LgGatC, MSP1E3D1, which is a membrane scaffold protein^21^, and *E. coli* lipids were combined at a molar ratio of 0.7:1:120. The mixture was incubated at 4°C for 1 h, and Bio-Beads SM2 (Bio-Rad) equilibrated with ultrapure water was added to remove the detergent. After gentle mixing at 4°C overnight, the sample was ultracentrifuged at 40,000 rpm for 30 min (Himac S55A2 rotor) and subsequently loaded onto a Superdex 200 10/300 GL column (GE Healthcare) preequilibrated with buffer (50 mM HEPES-NaOH (pH 7.0) and 150 mM NaCl) in the presence or absence of 10 mM D-xylose. The peak fractions were collected, concentrated using Amicon Ultra 50K NMWL (Merck Millipore), and stored at −80°C until use. The purification process under D-xylose-free conditions was performed using the same procedure as that for the D-xylose-containing conditions, with the exception of the substitution of D-xylose in the size exclusion chromatography buffer.

### Cryo-EM grid preparation

Quantifoil holey carbon grids (Cu R1.2/R1.3, 300 mesh) were glow-discharged at 7 Pa with 10 mA for 10 s using a JEC-3000FC sputter coater (JEOL) before sample application. A 3-μL aliquot of the nanodisc reconstituted with 4 mg/mL LgGatC was applied and blotted on the grids for 3 seconds at 100% humidity at 8°C, with a blot force of 10; the sample was then plunged into liquid ethane using a Vitrobot Mark IV (Thermo Fisher Scientific).

### Cryo-EM data acquisition and processing

Cryo-EM datasets were collected on a CRYO ARM 300 transmission electron microscope (JEOL Ltd., Japan), operating at an accelerating voltage of 300 kV, equipped with a cold-field emission gun, an in-column energy filter, and a direct electron detection camera (Gatan K3 Summit, Gatan Inc.) at SPring-8 (Hyogo, Japan). Movies were collected at a nominal magnification of ×60,000; this corresponded to a calibrated pixel size of 0.752 Å/pixel, with a total electron dose of 50 e^-^/Å^2^ subdivided into 50 frames via SerialEM^39^.

The Cryo-EM datasets were processed with CryoSPARC v4.5.3^40^. For the D-xylose condition dataset, where D-xylose was present, a total of 5,950 movies were aligned using patch motion correction, and the contrast transfer function (CTF) parameters were estimated using patch CTF estimation. The particles were automatically picked using Blob picker in CryoSPARC. The particles were subjected to two rounds of 2D classification, generating 2D class averages. Using these 2D class averages as templates, the particles were re-picked with the template picker from the 5,950 micrographs, resulting in 2,480,272 particles, and these particles were extracted with a downsampling via Fourier cropping to the pixel size of 3.008 Å. The particles were then subjected to 2D classification. Only ∼500,000 particles were subjected to Ab initio reconstruction (K=4), generating 3D initial model. The 2,480,272 particles were subjected to two rounds of heterogenous refinement (K=4, 5) to remove the junk particles. The best class containing 705,596 particles was re-extracted to 0.752 Å/pixel and refined using nonuniform (NU) refinement. These particles were subjected to CTF refinement and reference-based motion correction to generate a subset of 705,027 cleaned particles. These particles were subsequently subjected to no-aligned 3D classification and NU refinement imposing C2 symmetry, and a 3D cryo-EM map at an estimated overall resolution of 2.70 Å was attained. The local resolution was estimated by the Local Resolution Estimation job in CryoSPARC.

For the apo dataset, which refers to the condition without xylose, a total of 11,250 movies were aligned using patch motion correction, and the contrast transfer function (CTF) parameters were estimated using patch CTF estimation. The particles were automatically picked using blob picker in CryoSPARC. The particles were subjected to 2D classification, generating 2D class averages. Using these 2D class averages as templates, the particles were re-picked with the template picker form the 11,250 micrographs, resulting in 2,830,728 particles. The particles were then subjected to 2D classification. Only 1,505,724 particles were then subjected to Ab initio reconstruction and heterogeneous refinement (K=2, 6) to remove junk particles. The best class containing 264,815 particles was refined using nonuniform (NU) refinement (imposing C2 symmetry), and a 3D cryo-EM map at an estimated overall resolution of 3.27 Å was attained. The local resolution was estimated by the Local Resolution Estimation job in CryoSPARC.

### Model building and refinement of cryo-EM structures

An initial model of LgGatC was prepared using AlphaFold3^41^. The model was fitted into the EM density map in the presence of D-xylose using UCSF ChimeraX^42^, refined in real space using PHENIX^38^ with secondary structure and geometry constraints, and manually adjusted using Coot^37^ to better fit the EM density. The model was further refined in real space using PHENIX. The refined model in the presence of D-xylose was then fitted into the EM density map in the absence of D-xylose, modified, and refined in the same manner. The molecular models and cryo-EM maps were visualized using PyMOL (https://pymol.org) or UCSF ChimeraX. Model refinement and validation statistics are provided in Supplementary Table S2.

