## Supplemental Figures for "Structural basis of a phosphotransferase system xylose transporter"

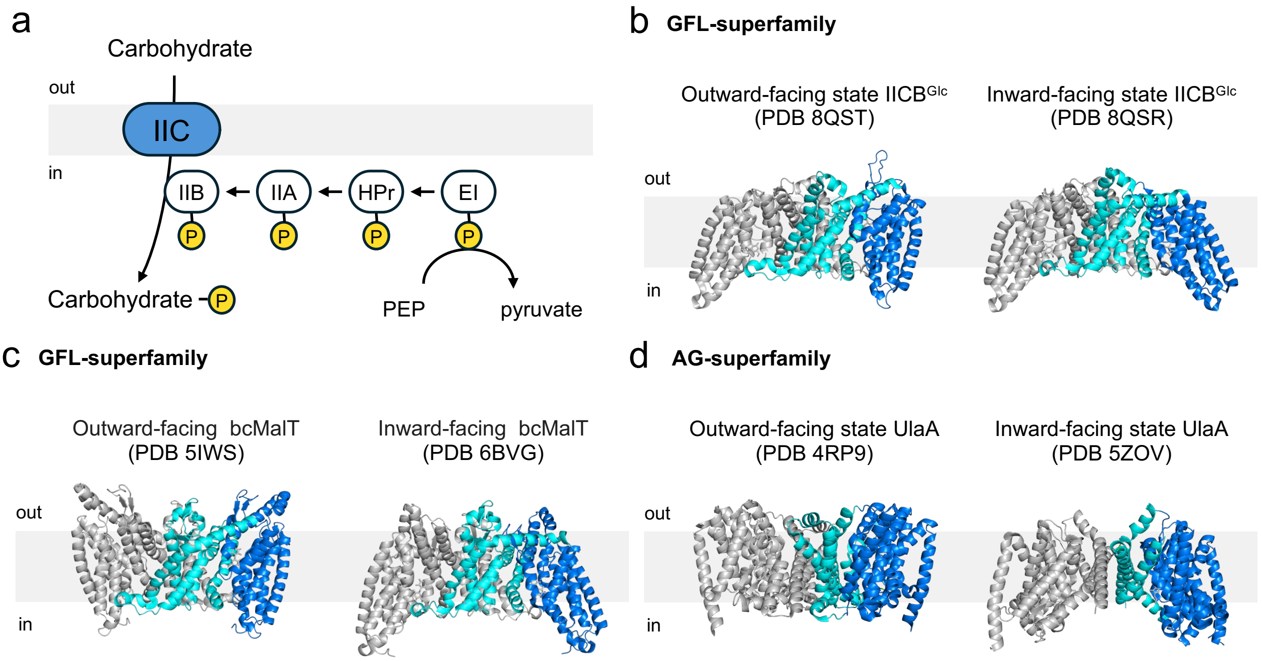


**Supplementary figure S1. Function and overall structures of PTS IIC**

(a) Schematic diagram of the carbohydrate uptake via the phosphoenolpyruvate-dependent phosphotransferase system (PTS). During or after uptake by the transporter IIC, the carbohydrate is phosphorylated by IIB, which forms a complex with IIC and receives a phosphate group sequentially transferred from PEP via EI, HPr, and IIA. (b–d) Structural comparisons of the outward- and inward-facing states of the following PTS IIC proteins: glucose-specific IICBGlc (b), maltose-specific IIC (bcMalT) (c), and L-ascorbate-specific IIC (UlaA) (d). IICBGlc and MalT belong to the GFL superfamily, whereas UlaA belongs to the AG superfamily. The interaction domains (scaffold or V motif domain) and a substrate-binding domain (transport or core domain) in a protomer are colored cyan and blue, respectively. The other protomers are colored gray. The PDB IDs are shown.

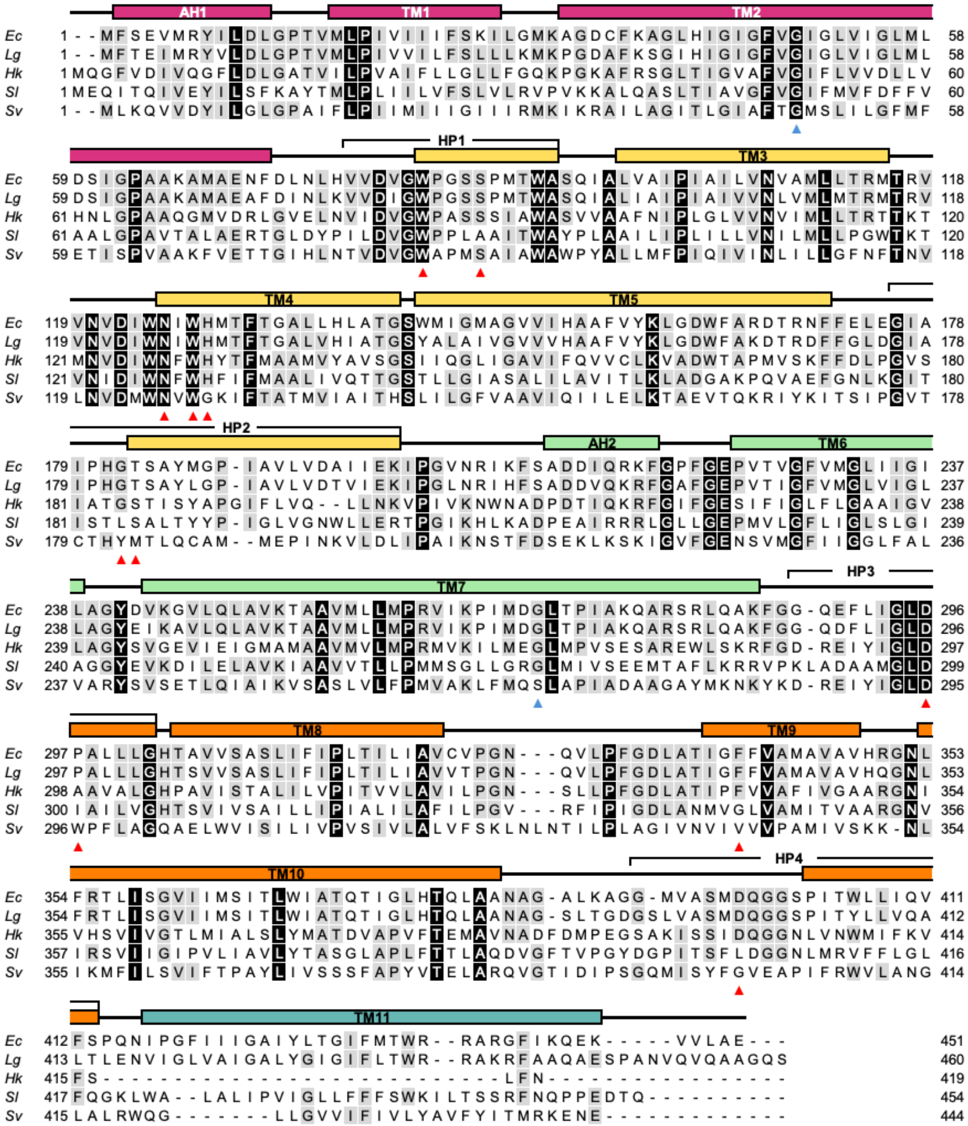


**Supplementary Figure S2. Multiple sequence alignment of GatCs.**

The alignment includes GatC orthologs from Escherichia coli (Ec) (UniProt: P69831), Leminorella grimontii (Lg) (UniProt: A0AAV5N9B3), Halobacillus karajensis (Hk) (UniProt: A0A024P4Y6), Spirochaeta lutea (Sl) (UniProt: A0A098QV03), and Sneathia vaginalis (Sv) (UniProt: A0A0E3UTQ6). Identical residues are highlighted in black. The α-helical regions of LgGatC are shown as bars above the alignment and are labeled with transmembrane regions (TM1–11), AH1–2, and HP1–4. The amino acid residues forming the cavity in LgGatC are indicated by red arrowheads, whereas the conserved glycine residues on TM2 and TM7 are indicated by blue arrowheads. The sequence identities of LgGatC with respect to EcGatC, HkGatC, SlGatC, and SvGatC are 81.37%, 41.05%, 31.93%, and 28.60%, respectively.

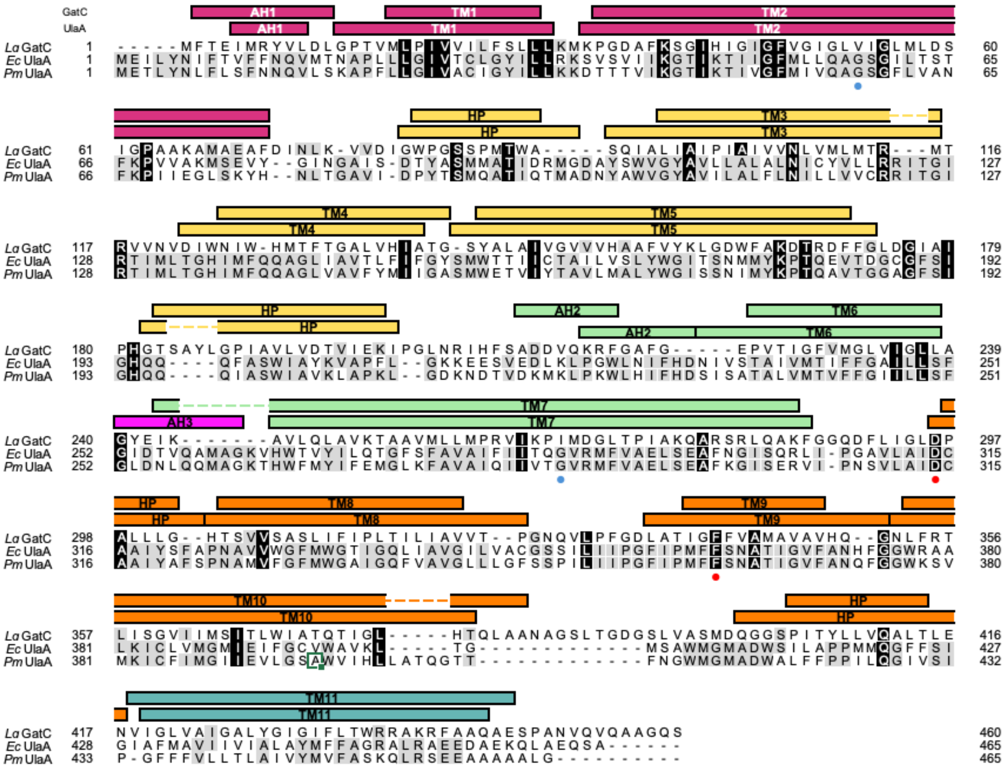


**Supplementary Figure S3. Amino acid residue sequence alignment of GatC and UlaA**

The alignment included Leminorella grimontii (Lg) GatC (UniProt: A0AAV5N9B3), Escherichia coli (Ec) UlaA (UniProt: P39301), and Pasteurella multocida (Pm) UlaA (UniProt: Q9CMQ1) and was manually prepared by superimposing and comparing their structures. The helical secondary structures of LgGatC and EcUlaA are shown above the sequence alignment. LgGatC shares sequence identities of 12.60% and 11.95% with EcUlaA and PmUlaA, respectively. Conserved amino acid residues located on the cavity surface are indicated by red circles. The glycine residues in UlaA that serve as the pivot points for the TM2 and TM7 curvatures are marked by blue circles. The AH between TM6 and TM7 of UlaA is shown in purple.

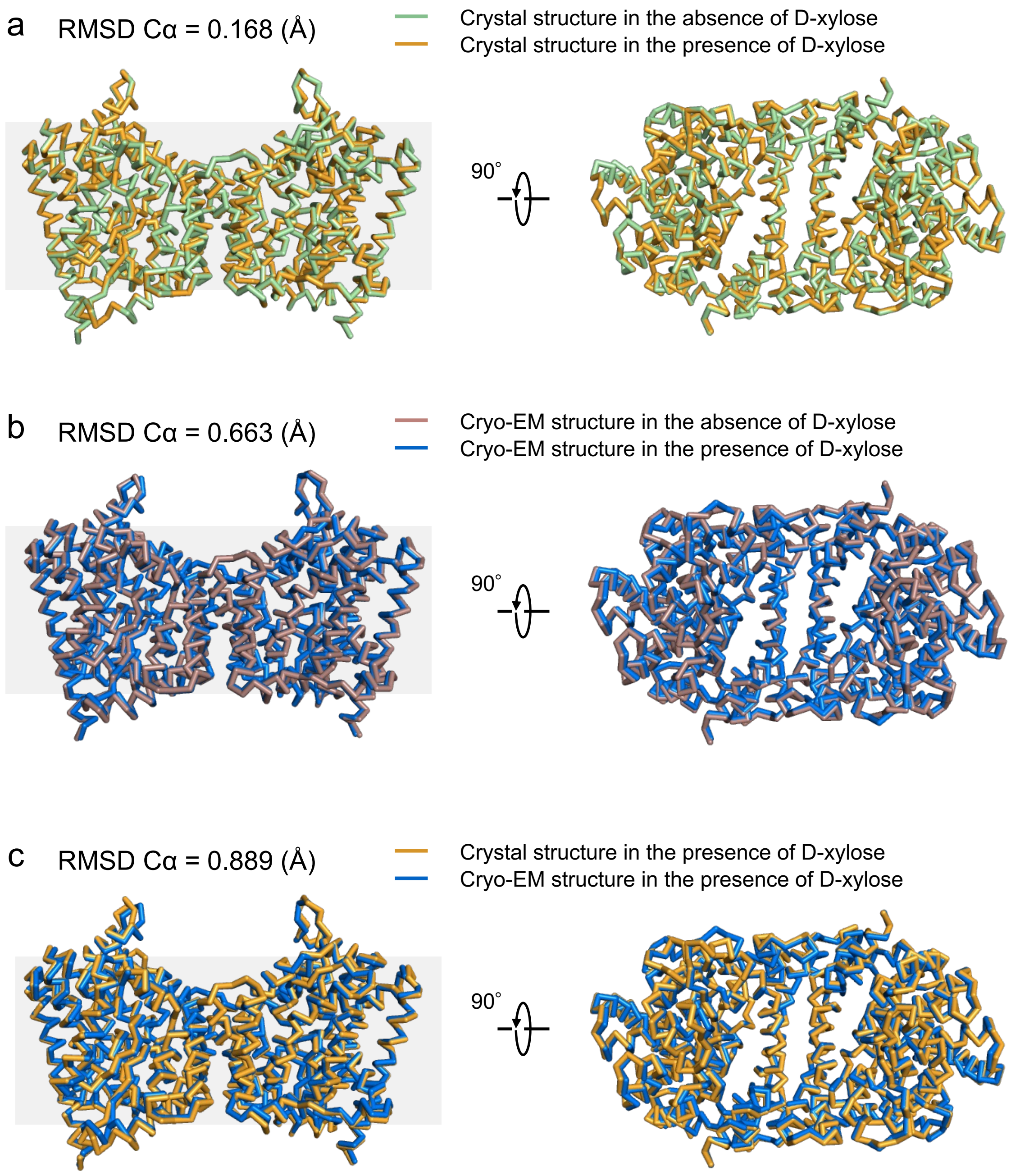


**Supplementary figure S4. Ribbon model of the crystal and cryo-EM structures of GatC with or without D-xylose.**
(a) The structural superposition of the crystal structures in the absence and presence of D-xylose. (b) Structural superposition of the cryo-EM structures in the absence and presence of D-xylose. (c) Structural superposition of the crystal and cryo-EM structures in the presence of D-xylose.

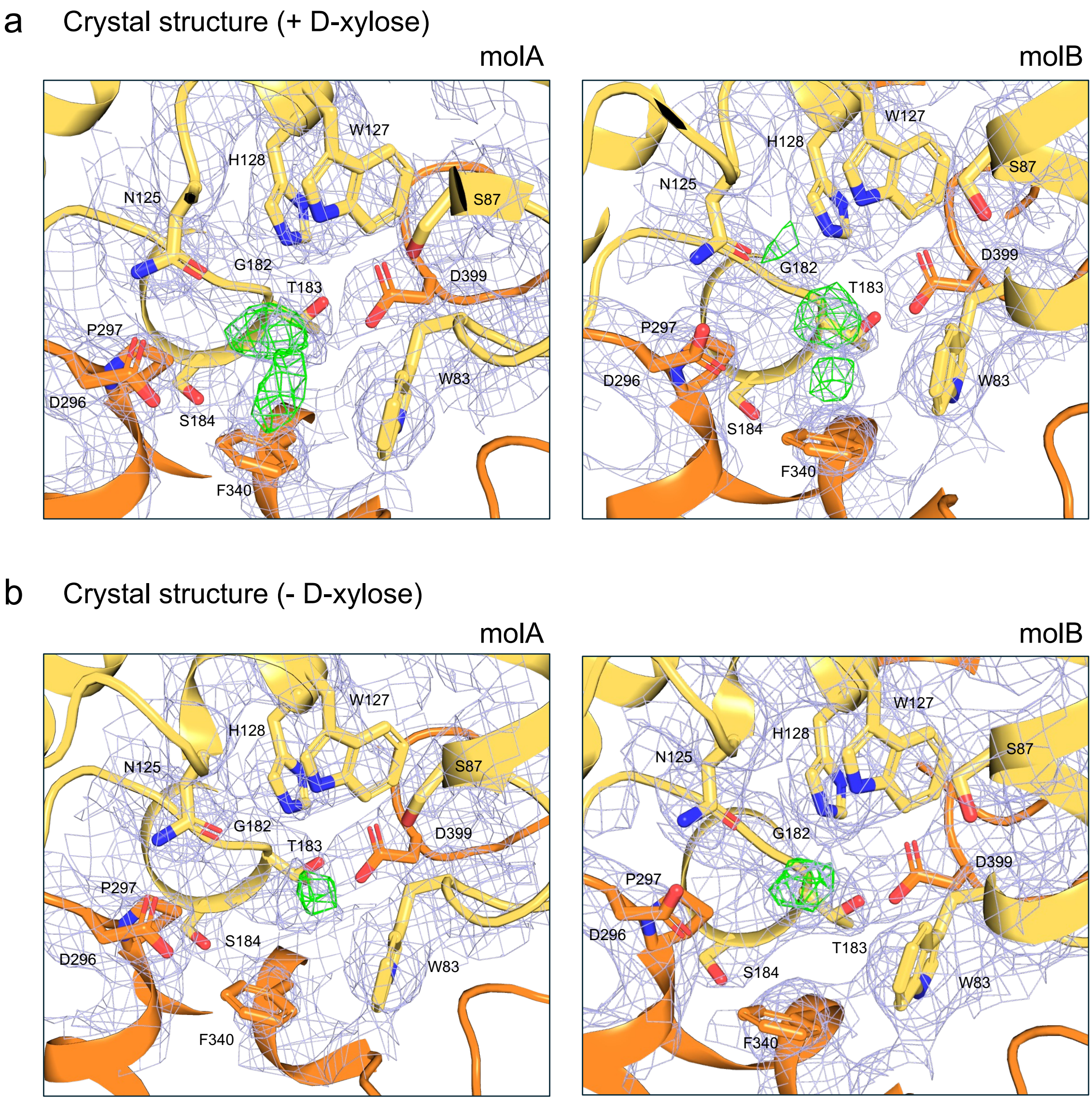


**Supplementary Figure S5. Structural comparison of D-xylose binding site in LgGatC crystal structures.**
Close-up view of the D-xylose binding site. (a, b) D-xylose binding site in the presence (a) and absence (b) of xylose. The 2Fo-Fc map and Fo-Fc map are shown with 1.0 σ (light blue) and 3.5 σ (green), respectively.

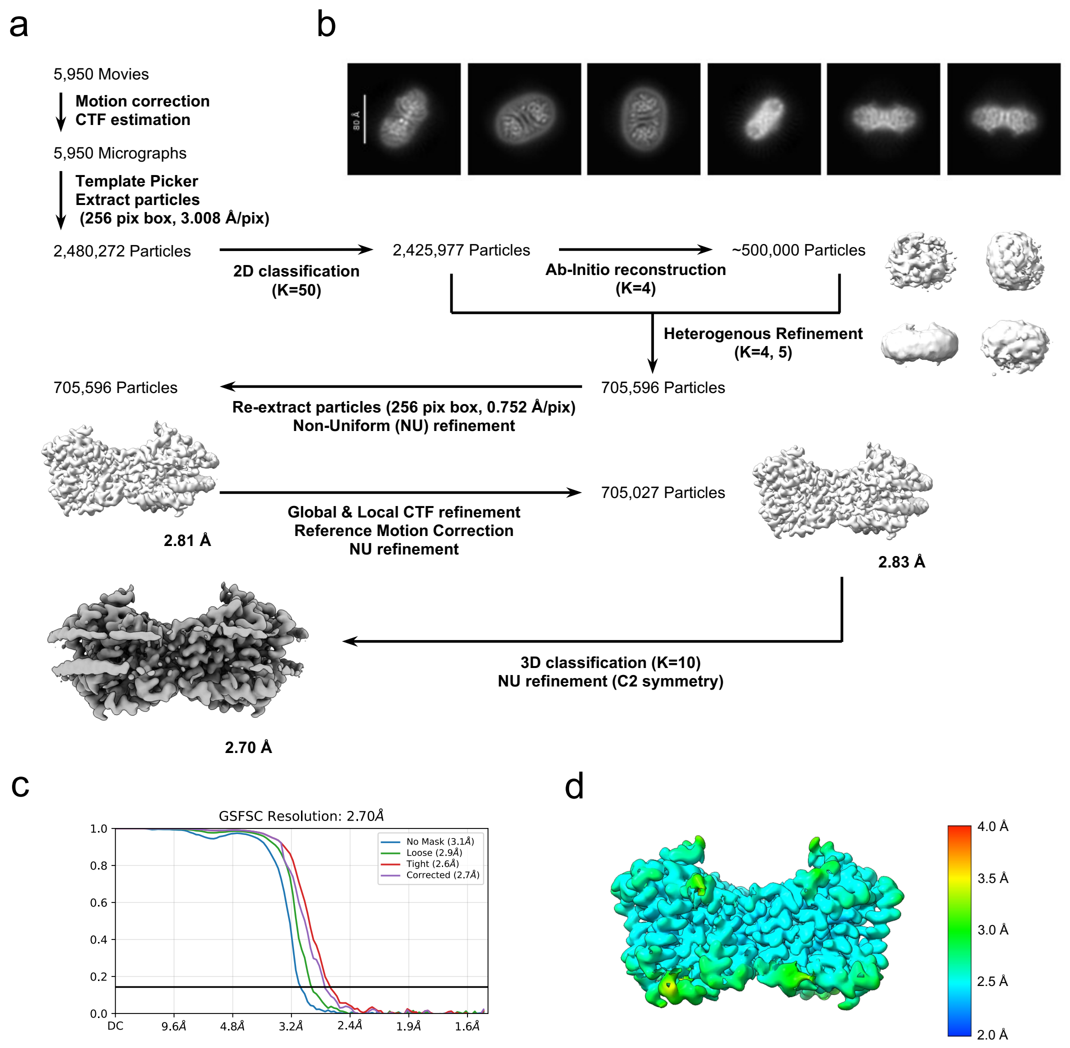


**Supplementary Figure S6. Cryo-EM analysis of the GatC in the presence of D-xylose.**
(a) Image processing workflow for LgGatC in the presence of D-xylose. (b) Representative 2D class averages. (c) Gold-standard FSC curve used for global-resolution estimates within CryoSPARC. (d) Final map colored according to the local resolution. The local resolution of the cryo-EM map was calculated using CryoSPARC.

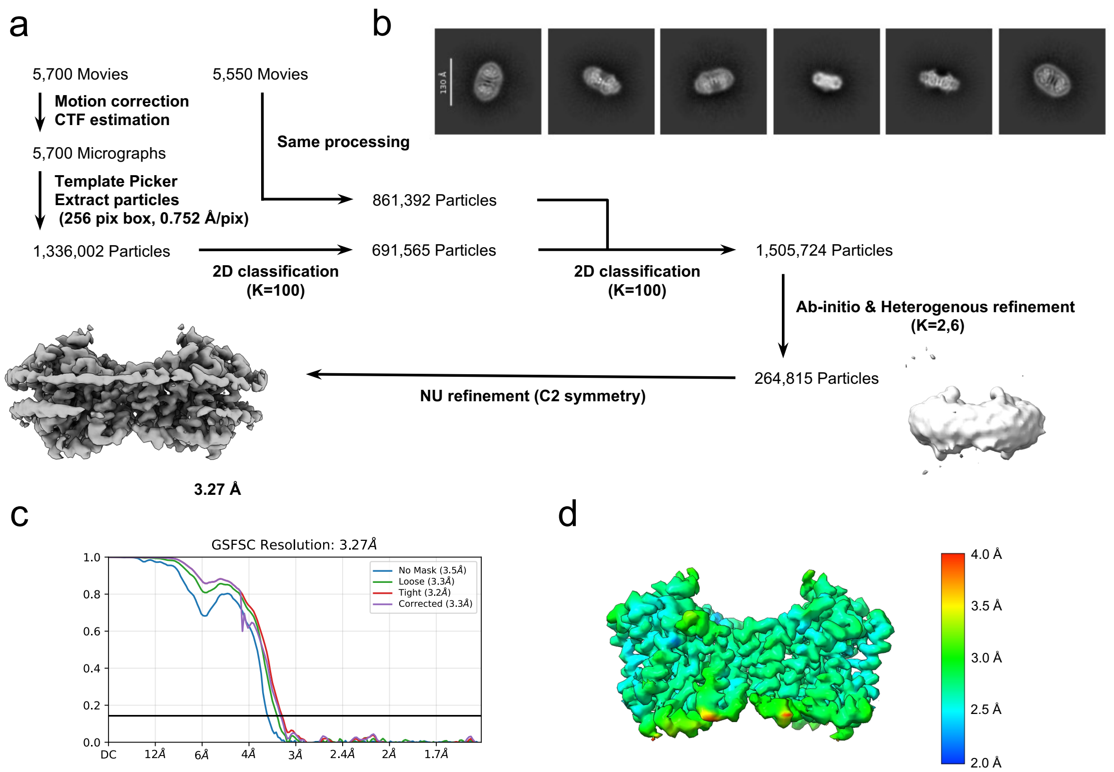


**Supplementary Figure S7. Cryo-EM analysis of the GatC in the absence of substrate**

(a) Image processing workflow for LgGatC in the absence of D-xylose. (b) Representative 2D class averages. (c) Gold-standard FSC curve used for global-resolution estimates within CryoSPARC. (d) Final map colored according to the local resolution. The local resolution of the cryo-EM map was calculated using CryoSPARC.

**Supplementary Table S1. X-ray diffraction data collection and refinement statics.**

| **PDB ID** | 9U8E | 9U8H |
|---|---|---|
| Crystal | **LgGatC in the presence of D-xylose** | **LgGatC in the absence of D-xylose** |
| **Data Collection** | | |
| X-ray source | SPring-8 BL32XU | SPring-8 BL32XU |
| Wavelength (Å) | 1.0 | 1.0 |
| Space group | $P22_12_1$ | $P22_12_1$ |
| A, b, c (Å) | 87.76 100.61 122.49 | 87.87 100.84 122.88 |
| Resolution range (Å) | 46.53 - 3.3 (3.4 - 3.3) | 46.65 - 3.74 (3.91 - 3.74) |
| Total reflections | 3157754 (250885) | 931086 (113807) |
| Multiplicity | 93.4 (88.7) | 41.6 (40.6) |
| Completeness (%) | 99.32 (99.93) | 99.12 (99.16) |
| Mean $I/\sigma$ ($I$) | 4.98 (0.43) | 3.03 (0.57) |
| Wilson B-factor | 53.90 | 61.25 |
| R-pim | 0.4203 (>5) | 0.4257 (>5) |
| $CC_{1/2}$ | 0.934 (0.394) | 0.859 (0.281) |
| **Refinement** | | |
| Reflections used in refinement | 16780 (1372) | 11679 (1409) |
| Reflections used for R-free | 1676 (137) | 1166 (141) |
| $R_{work}$ | 0.2223 (0.2691) | 0.2536 (0.3087) |
| $R_{free}$ | 0.2786 (0.3425) | 0.2855 (0.3449) |
| Number of non-hydrogen atoms | 6812 | 6812 |
| protein | 6708 | 6708 |
| monoolein | 104 | 104 |
| water | 0 | 0 |
| RMS derivation bond length (Å) | 0.004 | 0.004 |
| bond angles (˚) | 0.59 | 0.71 |
| Clashscore | 7.47 | 10.60 |
| Rotamer outliers (%) | 0 | 0 |
| Ramachandran favored (%) | 95.63 | 94.17 |
| allowed (%) | 4.37 | 5.83 |
| outliers (%) | 0 | 0 |
| Ramachandran outliers (%) | 0 | 0 |
| Average B-factor ($Å^2$) | 44.26 | 53.36 |
| protein ($Å^2$) | 44.17 | 53.43 |
| monoolein ($Å^2$) | 50.46 | 48.74 |
| water ($Å^2$) | - | - |

**Supplementary Table S2. CryoEM data collection and refinement statics.**

| | **LgGatC D-xylose-bound state** | **LgGatC Apo state** |
|---|---|---|
| **Data collection and processing** | | |
| EMDB entry | EMD-63950 | EMD-63951 |
| PDB entry | 9U82 | 9U84 |
| Microscope | CRYO ARM 300 | CRYO ARM 300 |
| Detecter | Gatan K3 summit camera | Gatan K3 summit camera |
| Magnification | 60,000x | 60,000x |
| Voltage (kV) | 300 | 300 |
| Data aquisiton software | SerialEM | SerialEM |
| Electron exposure (e $Å^{-2}$) | 49.92 | 50.32 |
| Defocus range (μm) | -1.4 to -1.6 | -1.4 to -1.6 |
| Pixel size (Å) | 0.752 | 0.752 |
| Symmetry | C2 | C2 |
| Micrographs | 5,950 | 11,250 |
| No. of initial particle images | 2,480,272 | 2,830,728 |
| No. of final particle images | 442,864 | 264,815 |
| Map resolution (Å) | 2.70 | 3.27 |
| FSC threshold | 0.143 | 0.143 |
| **Regfinement** | | |
| Initial model used | AlphaFold | 9U82 |
| Model resolution FSC threshold (Å) | 0.143 | 0.143 |
| Model resolution (Å) | 2.7 | 3.3 |
| Map sharpening B factor ($Å^2$) | 139.8 | 107.8 |
| Model composition | | |
| Non-hydrogen atoms | 6728 | 6708 |
| Protein | 896 | 896 |
| Ligands | XLS: 2 | 0 |
| B factors ($Å^2$) | | |
| Protein (min/max/mean) | 76.53/195.36/111.49 | 56.03/167.36/93.69 |
| Ligand | 107.93/114.31/112.02 | |
| R.m.s. deviations | | |
| Bonds lengths (Å) | 0.003 | 0.002 |
| Bond angles (°) | 0.534 | 0.538 |
| MolProbity score | 1.48 | 1.42 |
| Clashscore | 8.89 | 7.61 |
| Poor rotamers (%) | 0.71 | 0.85 |
| Ramachandran plot | | |
| Favored (%) | 99.55 | 98.65 |
| Allowed (%) | 0.45 | 1.35 |
| Outlier (%) | 0.00 | 0.00 |
